# Internal Tensile Force and A2 Domain Unfolding of von Willebrand Factor Multimers in Shear Flow

**DOI:** 10.1101/312405

**Authors:** Michael Morabito, Chuqiao Dong, Wei Wei, Xuanhong Cheng, Xiaohui F. Zhang, Alparslan Oztekin, Edmund Webb

## Abstract

Using Brownian molecular dynamics simulations, we examine the internal dynamics and biomechanical response of von Willebrand Factor (vWF) multimers subject to shear flow. The coarse grain multimer description employed here is based on a monomer model in which the A2 domain of vWF is explicitly represented by a non-linear elastic spring whose mechanical response was fit to experimental force/extension data from vWF monomers. This permits examination of the dynamic behavior of hydrodynamic forces acting on A2 domains as a function of shear rate and multimer length, as well as position of an A2 domain along the multimer contour. Force/position data reveal that collapsed multimers exhibit a force distribution with two peaks, one near each end of the chain; unraveled multimers, however, show a single peak in A2 domain force near the center of multimers. Guided further by experimental data, significant excursions of force acting on a domain are associated with an increasing probability for A2 domain unfolding. Our results suggest that the threshold shear rate required to induce A2 domain unfolding is inversely proportional to multimer length. By examining data for the duration and location of significant force excursions, convincing evidence is advanced that unfolding of A2 domains, and therefore scission of vWF multimers by the size-regulating blood enzyme ADAMTS13, happen preferentially near the center of unraveled multimers.

## INTRODUCTION

An ultra-large form of vWF (ULvWF) is synthesized and released into circulation by the Weibel-Palade bodies of endothelial cells containing as many as 3,500 monomers, and is essential for effective blood clot formation [1]. However, excessive concentrations of ULvWF can result in problematic blood clotting, so the enzyme ADAMTS13 cleaves ULvWF into a wide range of multimer lengths within about 2 hours of secretion [1]. Springer et al. reported that functional in vivo vWF multimers can contain up to 40 to 200 monomers [1]. Healthy hemostatic potential depends greatly on the proper size distribution and concentrations of vWF, which is dictated by the enzymatic cleavage of vWF by ADAMTS13 [2]. If vWF multimers do not undergo enough scission, then abnormally high concentrations of large vWF can cause the formation of many small blood clots called thrombi, leading to Thrombotic Thrombocytopenic Purpura (TTP) [3]–[5]. On the other hand, too much vWF multimer scission results in a lack of large vWF, which causes excessive bleeding and may lead to type I von Willebrand Disease (vWD) [6].

vWF assumes a globular conformation in normal circulation [7], [8]. However, large hydrodynamic forces can cause vWF to undergo macromolecular conformational changes by unraveling, often at the site of vascular damage [7]–[14]. In order to participate in necessary physiological processes, such as hemostasis, thrombosis, or scission, vWF must first be activated by a two-step process [1], [4], [15]; macromolecular unraveling is the first step towards activation. This work is primarily concerned with the second-step required for vWF scission activation. Although the underlying sensory mechanism remains unclear, it is generally accepted that the A2 domain senses elevated hydrodynamic force and unfolds accordingly [4], [10]. This mechanism is required for vWF cleavage to occur since the scission site is otherwise hidden in the folded A2 domain structure [14], [16]–[19]. The A2 domain remains folded under a threshold tensile force, making it unsusceptible to scission. Based on experimental results, Zhang predicted that the A2 domain unfolds around 11pN of tensile force [4]. For vWF in flow, achieving this unfolding force in an A2 domain greatly depends on macromolecular conformation, location along the multimer contour, and corresponding local internal tensile force distribution.

In addition to molecular scission, the platelet binding process also depends heavily on internal VWF dynamics, conformational activation, and tension dependent transitions. These things induce high affinity regions that enable local platelet binding [15]. Although the A2 domain is not directly responsible for vWF/platelet binding, a recent experimental work found that the A2 domain specifically binds to the active conformation of the A1 domain and effectively blocks interactions between the A1 domain and platelet GPIbα [20]. Another work employed both experiments and simulations in tandem and also reported specific A1/A2 domain binding that inhibits vWF/platelet binding; they found that an induced stretching force caused detachment of the two domains, thereby exposing the A1 domain binding site for GPIbα [21]. Clearly then, examination of internal vWF dynamics, namely A2 domain dynamics, is crucial for understanding vWF’s role in physiological processes at large.

For this reason, we have previously introduced a coarse-grained model that attempts to more realistically describe the molecular architecture inherent to vWF in order to better capture its internal dynamics, namely A2 domain dynamics [14]. While coarse-grained, our model achieves increased fidelity by explicitly modeling the A2 domain, which has been shown to unfold significantly under sufficient force [4]. The A2 domain has exhibited nonlinear force response to extension, although its real behavior is of course much more complex [4]. Nonetheless, its treatment as a FENE spring is sufficient for our purposes of examining internal force distribution in the current study. Our model of vWF has been parameterized on the monomer-level length scale, but more importantly our A2 domain represented FENE spring parameters have been parameterized according to experimentally obtained A2 domain unfolding data. A single vWF monomer is modeled by two beads connected by the parameterized A2 domain represented FENE spring, and adjacent monomers are connected by stiff harmonic springs, representing inter-monomer disulfide bonds.

Other authors have studied vWF cleavage experimentally and using BD simulations. Radtke et al. characterized the tension distribution along a vWF chain and employed a probabilistic condition to describe A2 domain susceptibility to cleavage, which derives from experimental findings by Lippok et al [9], [10]. Both authors were able to model vWF scission and predict a cleavage rate using a Morrison enzyme kinetics model. Huisman et al. introduced two vWF cleavage models, a geometric model and a force model, which utilize a cleavage probability based on the time step size and selected cleavage rate [22]. This allowed them to study the temporal evolution of vWF size distribution due to enzymatic cleavage for a given cleavage probability. Our work builds upon these foundational studies by modeling the A2 domain explicitly and deterministically assessing A2 domain susceptibility to cleavage by ADAMTS13 based on its internal force.

For several vWF multimer lengths, we directly calculate the shear flow conditions required to induce FENE spring force excursions that we associate with A2 domain unfolding. We claim that the most likely location for vWF scission by ADAMTS13 to occur is near to the multimer center by examination of internal tensile force and A2 domain unfolding duration distributions along the multimer contour.

## METHODS

*N* spherical beads of radius *a* are immersed in a fluid of viscosity *η*, which is exhibiting a simple shear flow given by 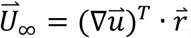, where 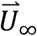 is the undisturbed velocity of a given bead purely due to the implicit solvent at location 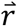 and 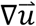 is the gradient of the pure solvent velocity. For a simple shear flow 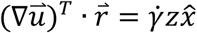, where 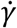 is the fluid shear rate and 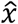 is a unit vector in the x-direction. Reynolds number based on the bead diameter is small so that Stokes flow is valid. Each bead is initially treated as a massless point force of friction, or drag, in the flow and induces a disturbance velocity. Thus, the velocity of the *i*^*th*^ bead is the sum of the pure solvent velocity and the *N* disturbance velocities at location 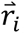. Note that a bead is subject to a drag force only when a velocity difference exists between the flow and the bead. Writing the Stokes equation for the *i*^*th*^ bead,

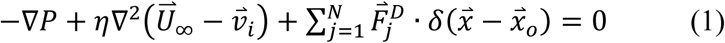

*P* is the pressure, 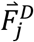 is the drag force on the *j*^*th*^ bead, and *δ* is the delta function which localizes the drag force to a point 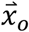. By the linearity of the Stokes equation, we find that the translational velocity of the *i*^*th*^ bead depends linearly upon the forces acting on it. That is,

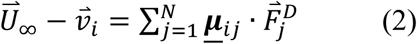

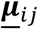 is the second-order mobility tensor, involving a Green’s function, between the *i*^*th*^ and *j*^*th*^ beads. Noting that 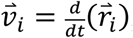, substituting the expression for the pure solvent velocity, and rearranging yields,

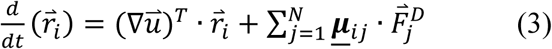

In the inertia-free limit, the drag force on any bead must equal the sum of the other forces acting upon it,

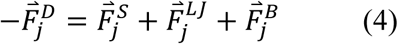

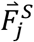 is the total spring force, 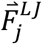 is the Lennard-Jones force, and 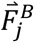 is the stochastic Brownian force. Substituting equation (4) into (3) and numerically integrating yields,

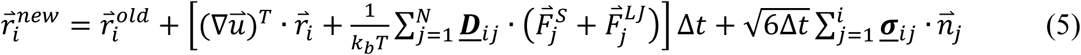

The second-order diffusion tensor is related to the mobility tensor by 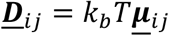, and describes Hydrodynamic Interactions (HI). 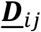 is given by the RPY approximation [23]–[25]:

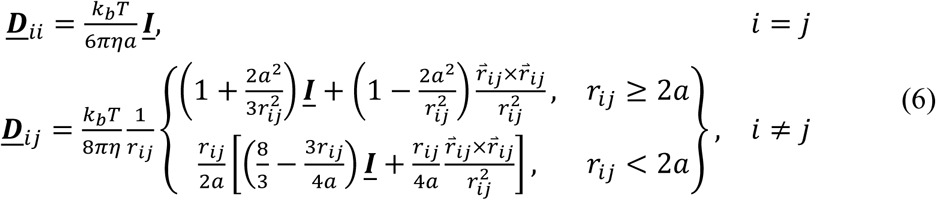

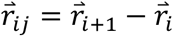 and 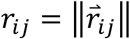. The selected solvent viscosity is *η* = 0.001 *pN μs/nm*^2^, which is the viscosity of water for a temperature *T* = 300*K* given in units physically germane to the problem. Kubo recognized one manifestation of the fluctuation-dissipation theorem relates bead mobility, or bead diffusion, to “fluctuation of the velocity of the Brownian motion [26].” Employing this relation in the treatment of random bead displacements requires calculation of the square-root of the fourth-order diffusion tensor, based on the assumption the random forces satisfy 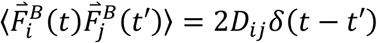 [27]. 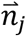 is a random vector with components uniformly distributed between [—1,1]. A factor of 3 is carried into the description because it is the magnitude of a random vector that is on average unit length. Ermak and McCammon used Cholesky decomposition to obtain the magnitude components of the random forces [28],

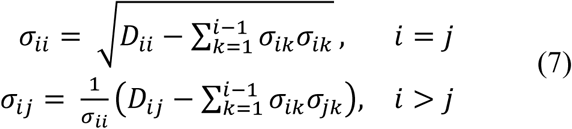

Quantities are made dimensionless by rescaling lengths by 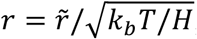 force by 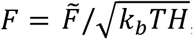 and time by 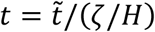. Beads are connected by either a Finitely Extensible Nonlinear Elastic (FENE) or harmonic spring. The spring forces are given by:

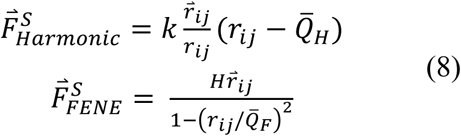

Experimental data from reference [4] was used to compute the FENE spring constant, *H* = 0.1428 *pN/nm*, and the spring length 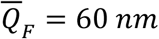 was chosen since the contour length of a vWF monomer is about 60nm [16]. The dimensionless harmonic spring force constant is *k* = 100, which was selected to model the strong inter-disulfide bonds between adjacent monomers [1], [16]. It oscillates around an equilibrium length of 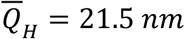, which was chosen to obtain an appropriate dimer length. Macromolecular cohesion is governed by the truncated 12-6 Lennard-Jones interaction force given by:

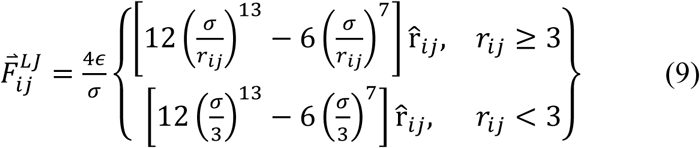

where *ϵ* and *σ* are the Lennard-Jones energy and length parameters, respectively. 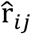 is the unit vector in the direction of 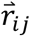. The selected bead radius is *a* = 10 *nm* and the length parameter is *σ* = 2*a*/2^1/6^ *nm*. The energy parameter is given by *ϵ* = 0.52 *k*_*b*_*T*, and was chosen to ensure that vWF multimers are in a collapsed state for the longest chain lengths herein considered.

Simulations began by initializing a vWF multimer of given length in a shear flow, where beads were placed either 4 or 5 dimensionless units apart depending on the connecting spring. Data were collected for 30τ, after the first τ was discarded to ensure no transient effects were considered in the averaged data. The initialization process is described in more detail in [14]. A multimer’s elongation relaxation time, *τ*, is based on the time required for an elongated chain to return to a globule [29]–[31], which increases with length and was determined previously in [14]. Multimer length is characterized by the value *M*, which indicates the number of monomers. Each monomer is comprised of two beads connected by a parameterized FENE spring. Results for a single simulation are averaged over 24 non-interacting chains, each of which possessed a unique random initial conformation and dynamic seed. Data collection occurred every 0.02*τ*, and the dimensionless time step chosen was Δ*t* = 10^−4^.

## RESULTS AND DISCUSSION

### a) Average Internal Tension

Figure 1 illustrates ensemble average Radius of Gyration (Rg) as a function of Weissenberg number (Wi) for all considered multimer lengths. Wi is defined as 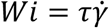, where τ is the polymer relaxation time and 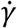 is the fluid shear rate. Rg increases with Wi, which becomes more pronounced past *Wi* = 1. Shorter chain lengths exhibit a gradual increase in Rg that is characteristic of coiled polymer elongation transitions, whereas longer chain lengths exhibit more dramatic increases in Rg that are indicative of the flow response for collapsed polymers – the behavior of coiled and collapsed polymers has been well characterized by prior authors [32], [33]. We can semi-quantitatively delineate the elongation behavior of shorter and longer chains upon the introduction of shear flow. Based on Fig. 1, it appears that chains of length 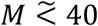 behave more like coiled polymers under the influence of shearing, whereas longer multimers exhibit collapsed polymer behavior. Other authors have also observed the dependence of Rg growth on chain length [32], [34]. For reasons revealed in a later discussion, we are primarily concerned with the dynamics of already unraveled or partially unraveled polymers in this work, which requires us to granulate simulation results according to the degree of polymeric elongation. This granulation reveals that the shape of the internal tensile force distribution does not depend on the coiled or collapsed nature of the polymer, but rather on the relative elongation attained by the polymer. Accordingly, the selected value of *ϵ* is sufficient for determining the internal tensile forces of unraveled polymers using the granulation techniques herein considered.

**Figure 1:**
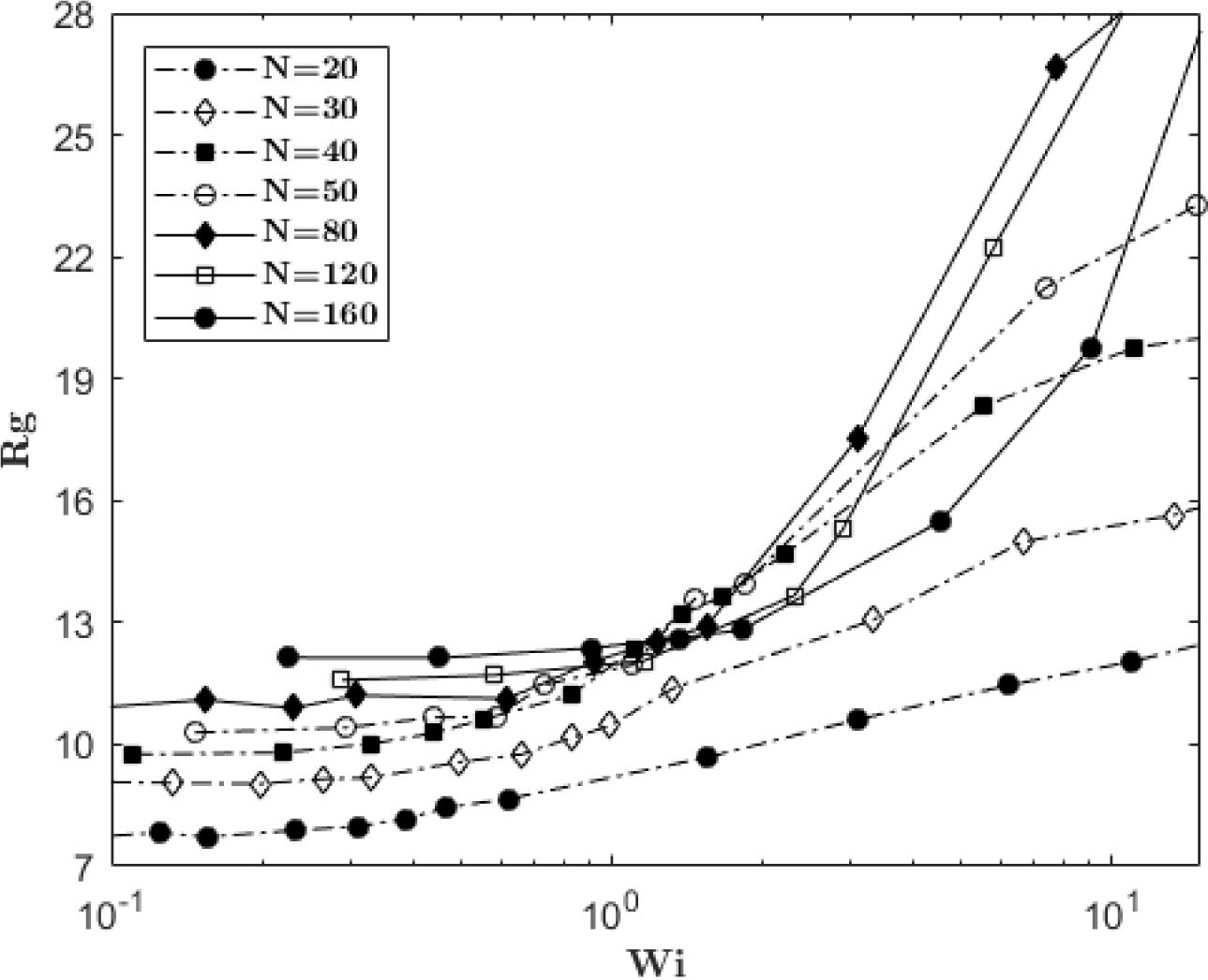
Dimensionless ensemble average radius of gyration for several vWF multimer lengths as a function of Weissenberg number. Line segments connecting individual datum indicate multimers behaving in a collapsed (solid segments) or coiled (dashed segments) manner.

Internal properties of vWF must be examined in order to understand its role in physiological processes. One such property is the internal tensile force in the A2 domain represented FENE springs. Fig. 2 illustrates ensemble average internal tensile force as a function of multimer length for various shear rates. Our interest in this work is to extend simulation considerations to in vivo conditions, accordingly we will hereafter present shear rates in dimensional units. Further, the magnitudes of the specific shear rates are obviously large and will be addressed later. Three different trends are illustrated in Fig. 2 that depend on the shearing strength. Tension is essentially uniform at zero flow and for the smallest shear rate illustrated, but actually, a very small, but statistically significant, monotonic decrease in force with increasing length is present. This is because macromolecular cohesion increases with multimer length due to increased Lennard-Jones attraction forces among beads, which increases the affinity for globule formation [32], [34]. For moderate shearing, force initially increases with length because drag forces exerted by the flow also increase, which must be internally balanced. However, internal tensile force stops monotonically increasing when multimer length becomes sufficiently long such that polymers become collapsed, at M ≅ 40. In other words, as multimer length increases so do hydrodynamic shielding and macromolecular cohesion that offer resistance to the unraveling tendencies of shear flow. For the largest shear rate illustrated in Fig. 2, shearing and hydrodynamic forces are so large that internal tensile force monotonically increases for all considered multimer lengths.

**Figure 2:**
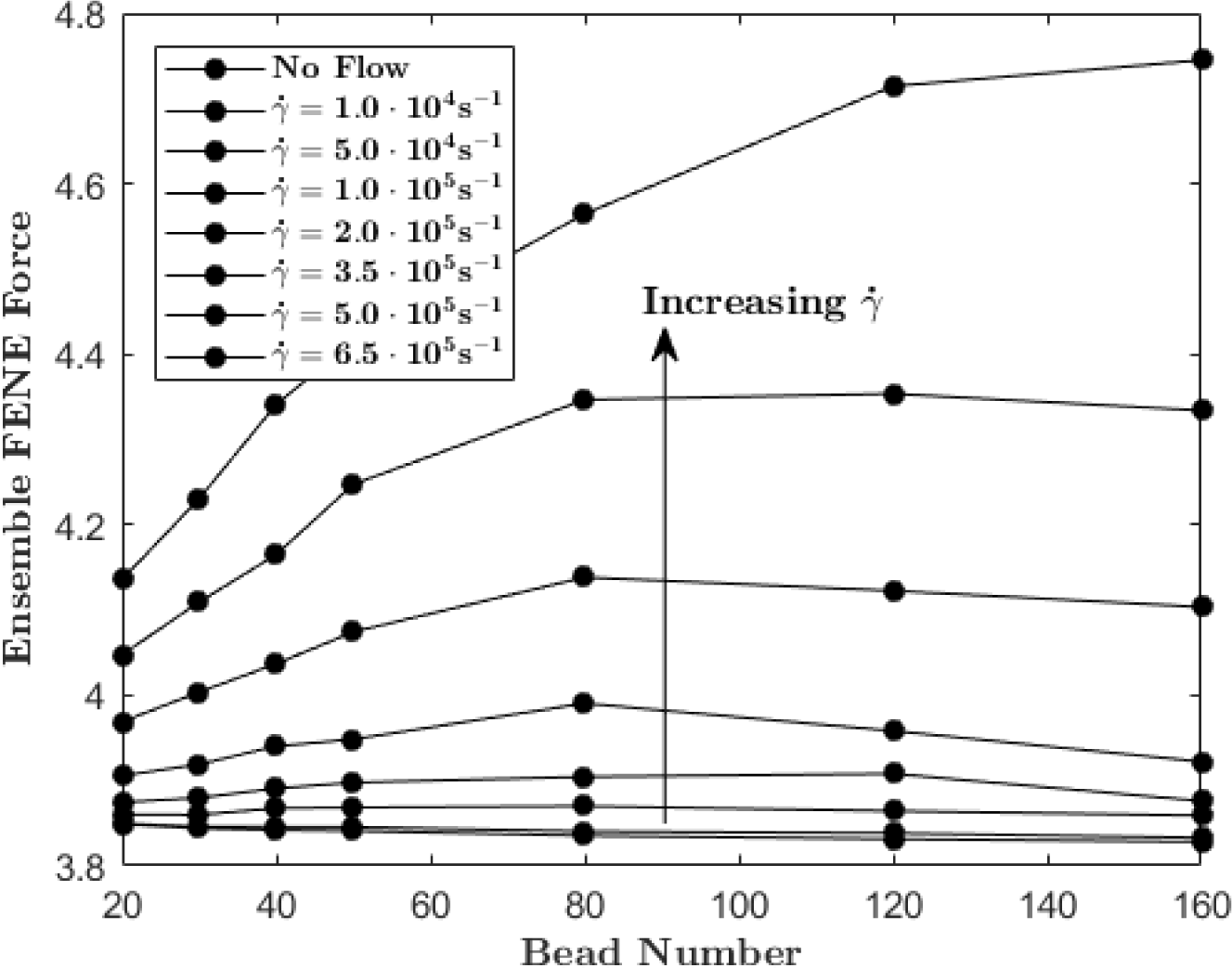
Dimensionless ensemble average internal tensile force within A2 domain represented FENE springs as a function of multimer length for several shear rates.

### b) Tension Distribution Dependency on Macromolecular Conformation

It is peculiar that ensemble average internal tensile force decreases with increasing chain length. This occurs after a certain length for most shear rates illustrated in Fig. 2. In fact, this result opposes what we intuitively expect to occur – that is, internal tensile force should monotonically increase with length due to elevated drag forces acting upon longer multimers that must be internally balanced. This is essential in the case of A2 domain unfolding because it is thought that internal tensile force is the activating mechanism and occurs around 11pN [4]. Trends in Fig. 2 wrongly suggest that longer chains are less likely to be cleaved, which is known to be false because ultra large vWF undergoes immediate scission upon secretion into the vasculature because of its length [1], [16]. However, the subtlety easily missed is that Fig. 2 illustrates the ensemble average internal force, which has been computed over all chain configurations during a simulation. In order to make sense of the trends in Fig. 2, we must examine the distribution of internal tensile force along the multimer contour.

Data in Fig. 3A represent average internal tensile force distributions at various stages of the macromolecular unraveling process for fixed multimer length and shear rate. At each data collection step for a given ensemble, molecules were binned according to the maximum distance between any two beads at that instant. Each bin represents 10% of the end-to-end distance for the perfectly aligned and relaxed multimer of corresponding length. The bottom distribution corresponds to all conformations throughout a simulation for which the maximum bead-bead separation distances are less than or equal to 10% of the perfectly aligned and relaxed chain length. Subsequent distributions represent the average only for conformations in the corresponding bin (10% - 20%, etc). The top curve represents conformations with the largest observed bead-bead separation distances, which were between 140% – 150% of the perfectly aligned and relaxed length.

**Figure 3:**
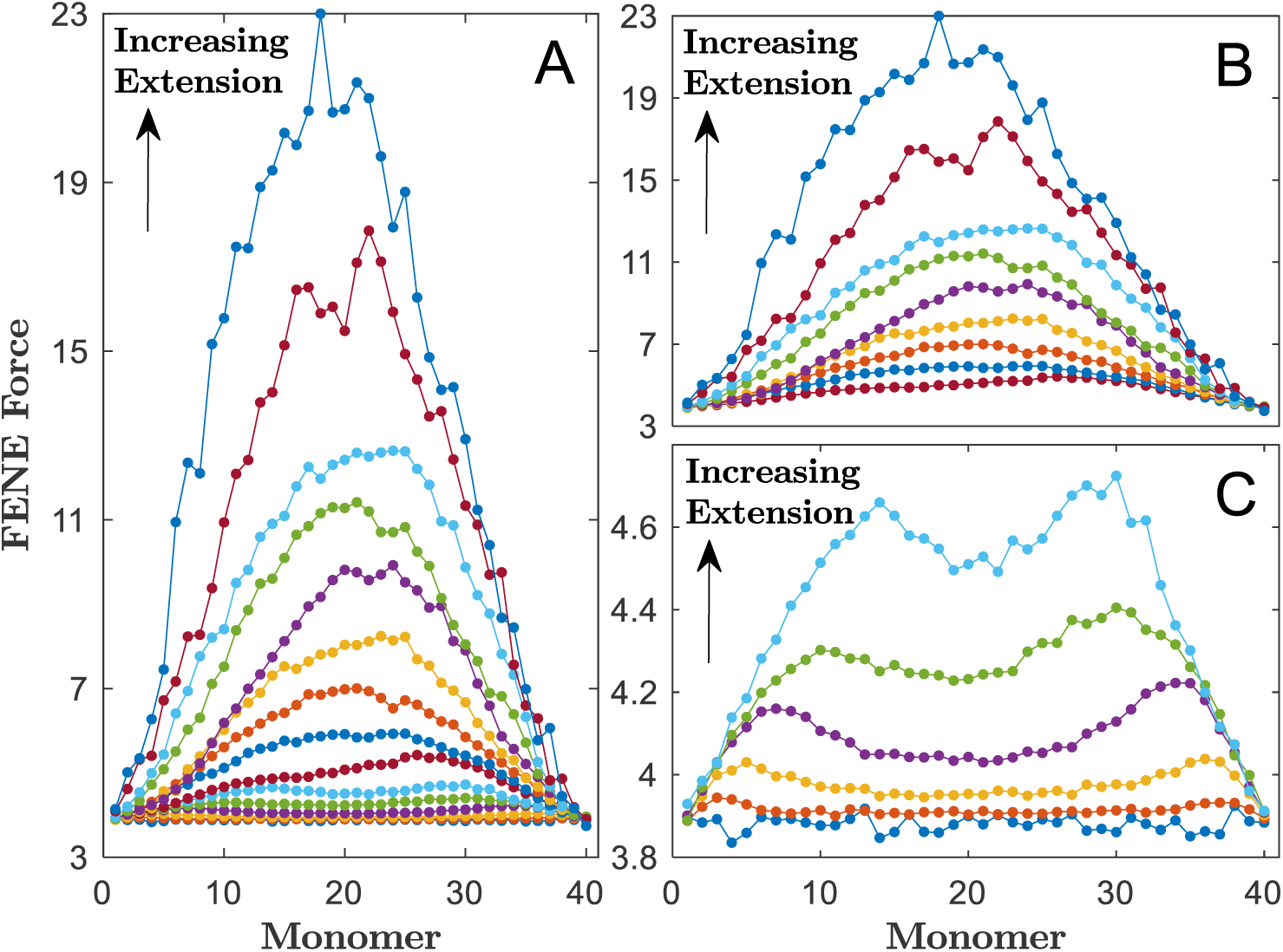
Dimensionless average internal tensile force distributions along the contour of a vWF multimer of length *N* = 80 (40-mer) at a shear rate of 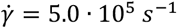. Increasing curves represent internal force distributions at various stages of the macromolecular unraveling process for (A) the entire process, (B) unraveled conformations, and (C) globular conformations.

Figure 3A exhibits two types of force distributions – associated with globular and unraveled macromolecular conformations. Similar to what other authors have reported, internal tensile force distributions for globular conformations exhibit a double-peak structure, where the peaks are located near the chain termini [9], [10]. The unraveling process is rendered in Fig. 4, and usually begins when a multimer end escapes from the globule (i.e. a protrusion nucleates) [9], [10], [32], [34]. Internal force increases in the protruding strand because it is no longer protected from flow by hydrodynamic shielding. As macromolecular unraveling continues, force increases at the double-peak structures and the peaks move nearer to the multimer center – indicating longer protrusions from the globule. Internal force distributions for globules of increasing protruding strand length are illustrated in Fig 3C. Eventually, the hydrodynamic force on the protrusions becomes so large that the globule unravels, and the double-peak structures merge into one peak located at the multimer center. This characterizes the internal tensile force distribution for unraveled vWF, and by inspection of Fig. 3A and Fig. 3B, the shape is roughly proportional to the square of length.

Figure 3A illustrates force distributions for the complete unraveling process and they have been decomposed according to macromolecular conformation in the adjacent subfigures. Fig. 3B and Fig. 3C illustrate the internal force distributions for unraveled and globular conformations only, respectively. The transition from a globular to unraveled conformation occurs when the internal tensile force distribution changes from a double-peaked structure to parabolic. For all chain lengths herein considered, this transition occurred when the maximum distance between any two comprising beads was ~60% of the end-to-end distance for the perfectly aligned and relaxed multimer of the corresponding length. Determining this transition percentage for smaller shear rate becomes challenged by the rare event nature of both molecular and domain unfolding, along with the concomitant lack of data on unfolded conformations.

**Figure 4:**
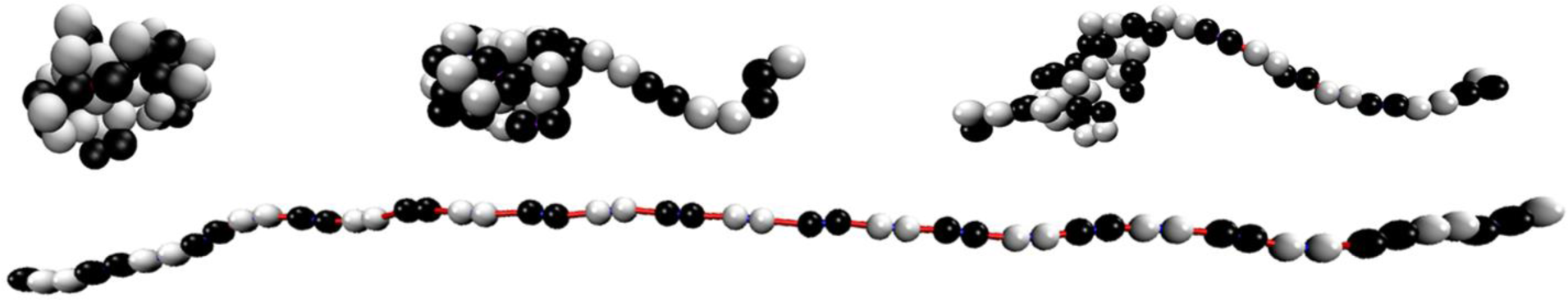
A vWF chain of length *N* = 50 beads (25-mer) at various stages of the macromolecular unraveling process [35]. A monomer is represented by two beads, one black and one white, connected by a red FENE spring. Short, stiff harmonic springs connect beads of like color. They represent strong inter-disulfide bonds connecting adjacent monomers in a head-to-head and tail-to-tail formation. Individual monomers are well shielded from flow in compact globular conformations. Macromolecular unraveling usually begins at the multimer termini in bulk flow. Note that the centermost A2 domain represented FENE springs are significantly unfolded in the fully unraveled conformation, whereas springs near the base of the longest protruding strand are only slightly stretched.

Returning to the dilemma posed in the discussion of Fig. 2, we wish to address why ensemble average force decreases with increasing length for certain chain lengths and shear rates. From Fig. 3, we observed that two distinct types of force distributions exist during the unraveling process, corresponding to globular and unraveled conformations. Further, we learned that this transition occurs for extensions that are ~60% of the maximum but relaxed end-to-end length. It is natural then to further granulate simulation results according to these two conformations, illustrated in Fig. 5 and Fig. 6.

**Figure 5:**
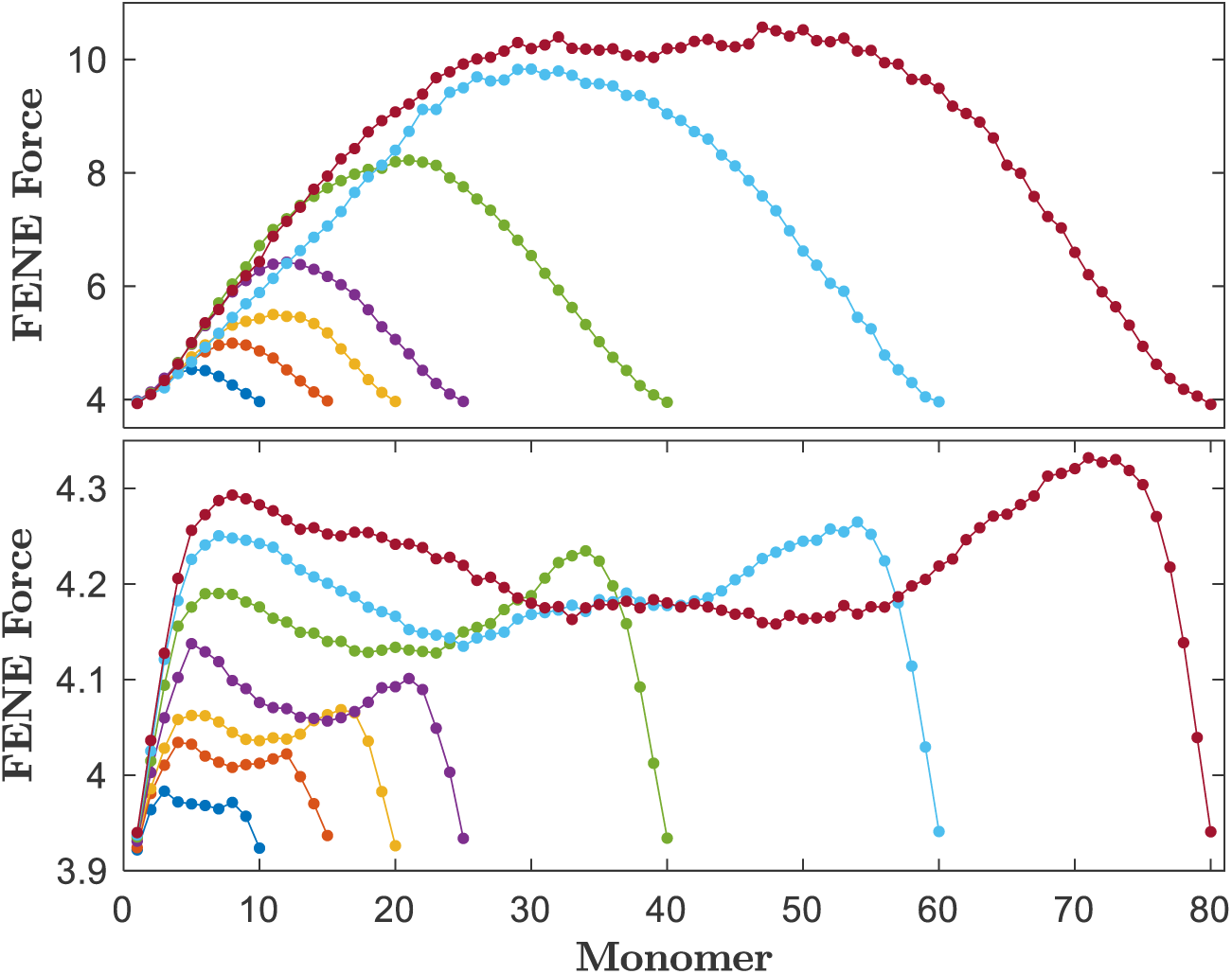
Dimensionless average internal tensile force distributions for various multimer lengths at a shear rate *γ* =3.5· 10^5^ *s*^−1^ as a function of monomer position for (top) unraveled and (bottom) globular conformations only, respectively.

**Figure 6:**
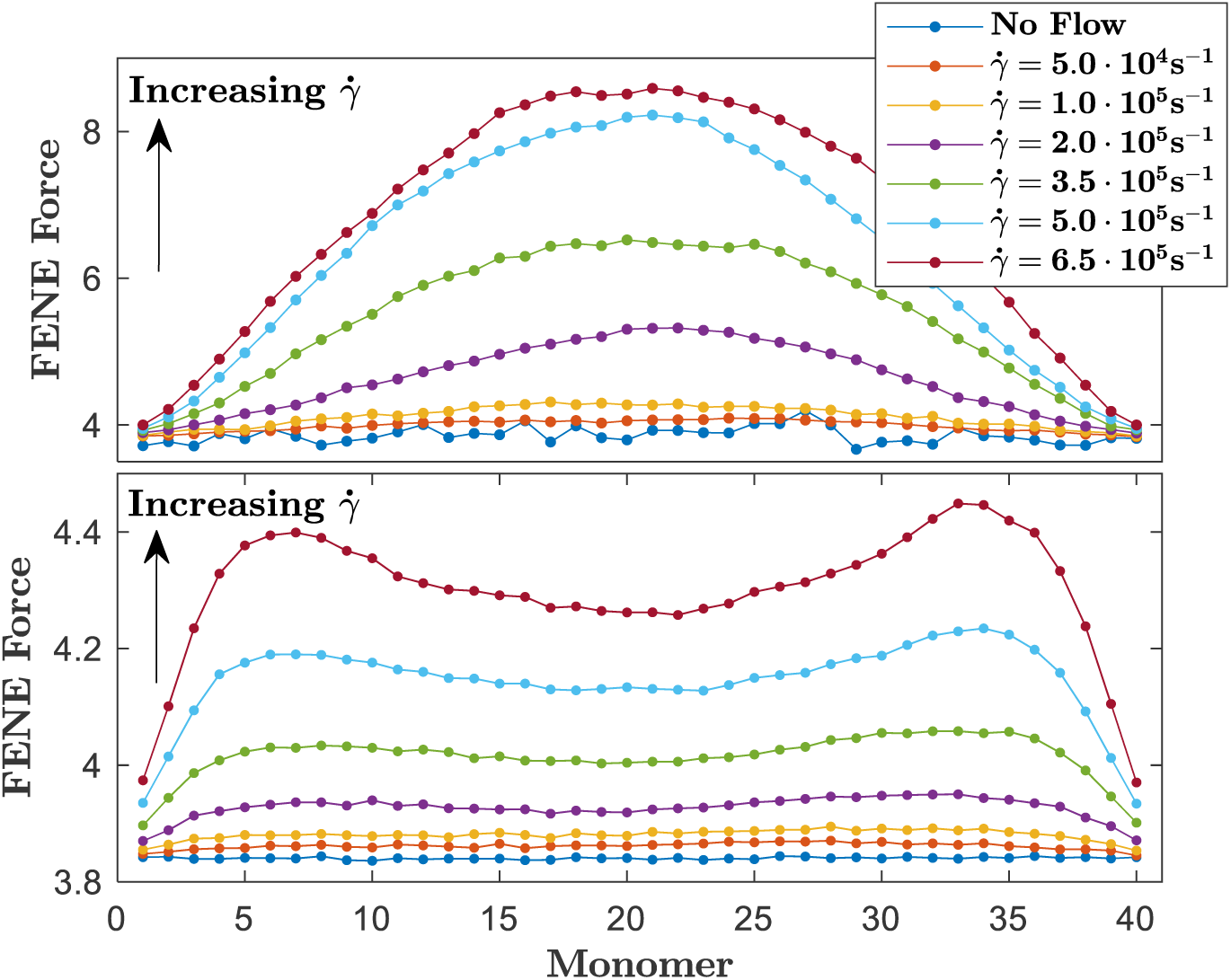
Dimensionless average internal tensile force distributions for a multimer of length *N =* 80 (40-mer) as a function of monomer position at several shear rates for (top) unraveled and (bottom) globular conformations only, respectively.

Figure 5 illustrates internal tensile force distributions along the multimer contour at a shear rate 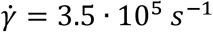 for unraveled and globular conformations only, respectively. A multimer was considered unraveled (or globular) if the largest bead-bead separation distance was at least (or at most) 60% of the end-to-end distance for the corresponding perfectly aligned and relaxed length. Fig. 5 shows that further granulation of force data according to macromolecular conformation yielded the anticipated trends between force and chain length – that is, internal tensile force monotonically increases with length. Therefore, the local internal tensile force distribution is heavily dependent upon macromolecular configuration. Fig. 5 shows that internal tensile force monotonically increases with chain length only when hydrodynamic forces are actively unraveling the multimer. Internal tensile force magnitudes are much smaller for globules than unraveled conformations due to hydrodynamic shielding. Fig. 6 illustrates internal tensile force distributions for a vWF chain of length M = 40 at several shear rates for unraveled and globular conformations only, respectively. For small shear rates, globular conformation force distributions are nearly homogeneous. The drastic differences in force magnitudes between globular and elongated conformations, illustrated in Fig. 5 and Fig. 6, indicate that ensemble average data in Fig. 2 are dominated by contributions from globular configurations (i.e. chains are globular more often than not). However, force monotonically increases with chain length for the largest shear rate considered in Fig. 2, which indicates A2 domain unfolding frequency has increased sufficiently that the ensemble averages become dominated by unraveled configuration contributions.

### c) Internal Tension and A2 Domain Unfolding

In order to examine A2 domain kinetics, we define an A2 domain unfolding event as any force excursion past the dimensional force value of 11 *pN* for some duration of time; we will discuss the durations of force excursions later. Unraveled conformations attain much higher internal tensile forces in flow, which suggests they are more likely to facilitate A2 domain unfolding than globules. Further, the shape of the distribution suggests that unraveled vWF chains are most susceptible to cleavage by ADAMTS13 at the multimer center, where internal force is greatest and the potential of A2 domain unfolding is highest. In that case, molecular scission favors a fractal-like behavior and may offer some explanation to the discrete and exponential size distribution exhibited by in vivo vWF concentrations [36]. Radtke et al. advanced that it might be possible for cleavage to occur at sites of maximal tensile force; however they also pointed out that it remained unclear to what extent the distribution of force influenced the cleavage process [10].

The high shear rates explored here bear discussion. Our monomer model uses two beads, each representing a collection of domains on either side of the A2 domain; the size of these beads is based on the experimentally observed size of vWF monomers [37], [38]. As a result, the value of bead radius employed here (*a* = 10 *nm*) is much lower than what has been used by a number of prior authors who described the vWF monomer or dimer with a single bead (*a* = 30 to 250 *nm*) [9], [10], [22], [39], [40]. This means hydrodynamic force applied on beads at a given shear rate will also be much lower. Further, the FENE spring used to represent the A2 domain has no size with regard to hydrodynamic interactions – that is, we neglect hydrodynamic drag acting on the A2 domain that tethers two beads and do not consider it when computing HI. This force is negligible for large bead radii, but becomes more relevant for the size of bead considered here, especially when the A2 domain begins to elongate. Perhaps the most notable fact is that we desire to observe what - in physiological blood flow regimes - should be a rare event. That is, many prior authors have advanced that the rate-limiting step for vWF scission by ADAMTS13 is the unfolding of an A2 domain [9], [10]. The concentration of the enzyme and the corresponding diffusion time (and reaction time) are considered secondary to the unfolding event. As such, it has been previously advanced by various groups that A2 domain unfolding should be associated with a scission event. Even for ultra-large vWF molecules secreted into blood, it takes time scales of order minutes to hours for the molecular size distribution to be reduced to physiological functional ranges [1]; thus, unfolding events - even for such large molecules - happen at a relatively low frequency. For vWF molecules in the functional size range, it is expected that A2 domain unfolding (and concomitant scission) is very rare in typical blood flow. The total duration of the longest simulations presented here is *27.2 ms;* furthermore, our largest multimers studied are still well below what is believed to be the upper limit for the functional size range. Thus, for physiological flow rates, we expect our model to exhibit zero significant force excursions over the duration of the simulation. Instead, we seek to observe many such events so that we can build a statistical description of them. Studying very large molecules or significantly longer simulation durations is computationally intractable; thus, we instead increase the driving force for unfolding (i.e. the shear rate) to ranges where many force excursion events can be observed, even for the smallest chains studied here. In discussing results below, the influence of the high shear rates on observations made is further addressed.

We wish to use our vWF model to examine the chain length dependence of the shear rates required to induce A2 domain unfolding in the bulk. We previously stated that A2 domain unfolding is most likely to occur for unraveled multimers near to the chain center. This is what we were referring to earlier in the discussion of Fig. 1 when we said that our primary concern in this work is with unraveled or partially unraveled multimers. Since we have explicitly modeled the A2 domain as a FENE spring, we can directly examine A2 domain unfolding kinetics. Zhang et al determined the A2 domain unfolds at an internal tensile force around 11pN [4], which corresponds to the dimensionless FENE spring force of 14.3. We consider an A2 domain represented FENE spring to be unfolded and susceptible to scission by ADAMTS13 for FENE forces greater than or equal to this threshold value.

Figure 7 illustrates average peak internal tensile force for unraveled conformations as a function of shear rate for all considered chain lengths. Data for a given ensemble were obtained by first computing the maximum internal tensile force in any A2 domain represented FENE spring along the multimer contour at each data collection step, then we computed the average value only for chains in unraveled conformations. A chain was considered unraveled if the largest bead-bead separation distance was at least 60% of the end-to-end distance for the perfectly aligned and relaxed multimer of corresponding length. Next, we normalized the average force value by the threshold dimensionless unfolding force 14.3, which corresponds to 11pN. We repeated this procedure for all chain lengths and shear rates and plotted the resulting averages. Force data for all chain lengths in Fig. 7 were well fit by a linear relation, 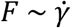.

**Figure 7:**
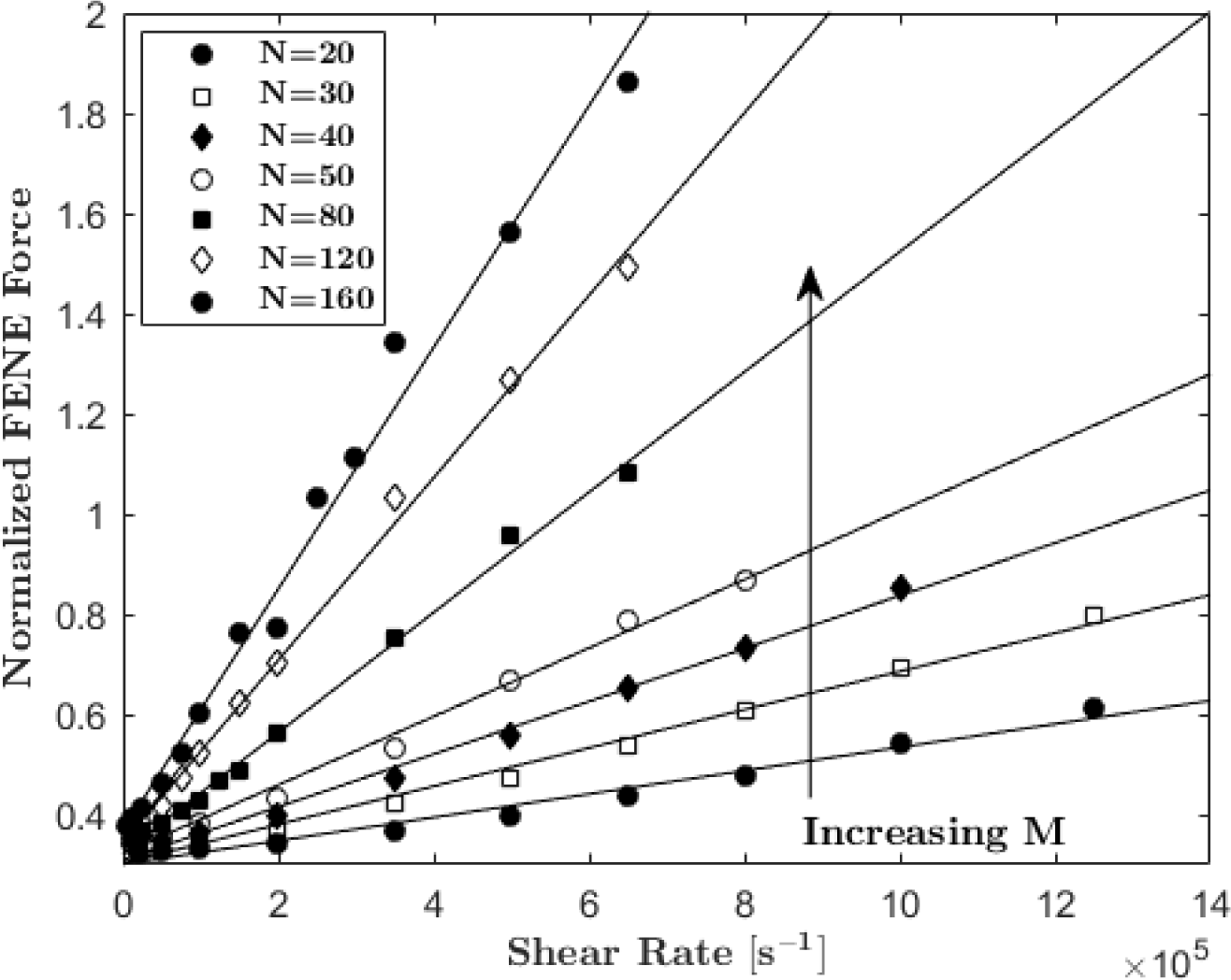
Normalized average peak internal tensile force for unraveled conformations as a function of shear rate for several multimer lengths. Peak force is normalized by the experimentally reported threshold dimensionless A2 domain unfolding force that corresponds to 11 *pN* [4]. The black curves indicate force data is well fit by a linear function of shear rate for all multimer lengths.

Internal force distributions illustrated in Fig. 5 and Fig. 6 indicate that peak force values in Fig. 7 correspond, on average, to the force present at the multimer center of unraveled chains, which is where vWF is most susceptible to cleavage by ADAMTS13 based on force induced A2 domain unfolding. Employing the macromolecular extension threshold criteria is justified because A2 domain unfolding occurs very infrequently, or not at all, for globules because they lack the internal force required to unfold the A2 domain due to shielding and cohesion effects. This phenomenon prevents excessive scission of vWF in the physical system. Further, the specific value of 60% is uniform among the considered chain lengths as the extension threshold for which globules transition into unraveled conformations, as discussed previously in regards to Fig. 3. The normalized value of one indicates that, on average, the centermost A2 domain is unfolded due to sufficient internal tensile force at the corresponding shear rate, and it is therefore susceptible to cleavage by ADAMTS13. We used the curve fittings to extrapolate or interpolate the normalized force value of one to obtain the corresponding shear rate.

The middle data set and accompanying curve fit in Fig. 8 illustrate the shear rates corresponding to normalized force values of one obtained from the curve fittings illustrated in Fig. 7 as a function of chain length. The curves in Fig. 8 relate multimer length to the critical shear rate required to unfold at least the centermost A2 domain represented FENE spring for an unraveled vWF chain. It has been assumed here that, if a single A2 domain (i.e. one that is experiencing the peak force near the center of an unraveled molecule) is subject to a force of 11 *pN*, then it will unfold. Experimentally however, unfolding was observed over a distribution of pulling forces that depended upon pulling rate; authors reported the most likely unfolding force for each pull rate, with values ranging from 7 *pN* up to 15 *pN* [4]. Obviously, unfolding is a complex molecular process that cannot be associated with a single force value; furthermore, it is unknown if unfolding in vivo differs from in the experimental environment. Given the complexity of the dependence of unfolding on applied force, for the analysis here, we have illustrated the dependency for the reported most likely unfolding force of 11 *pN* as well as the boundary values of experimental unfolding events of 7 *pN* and 15 *pN*.

**Figure 8:**
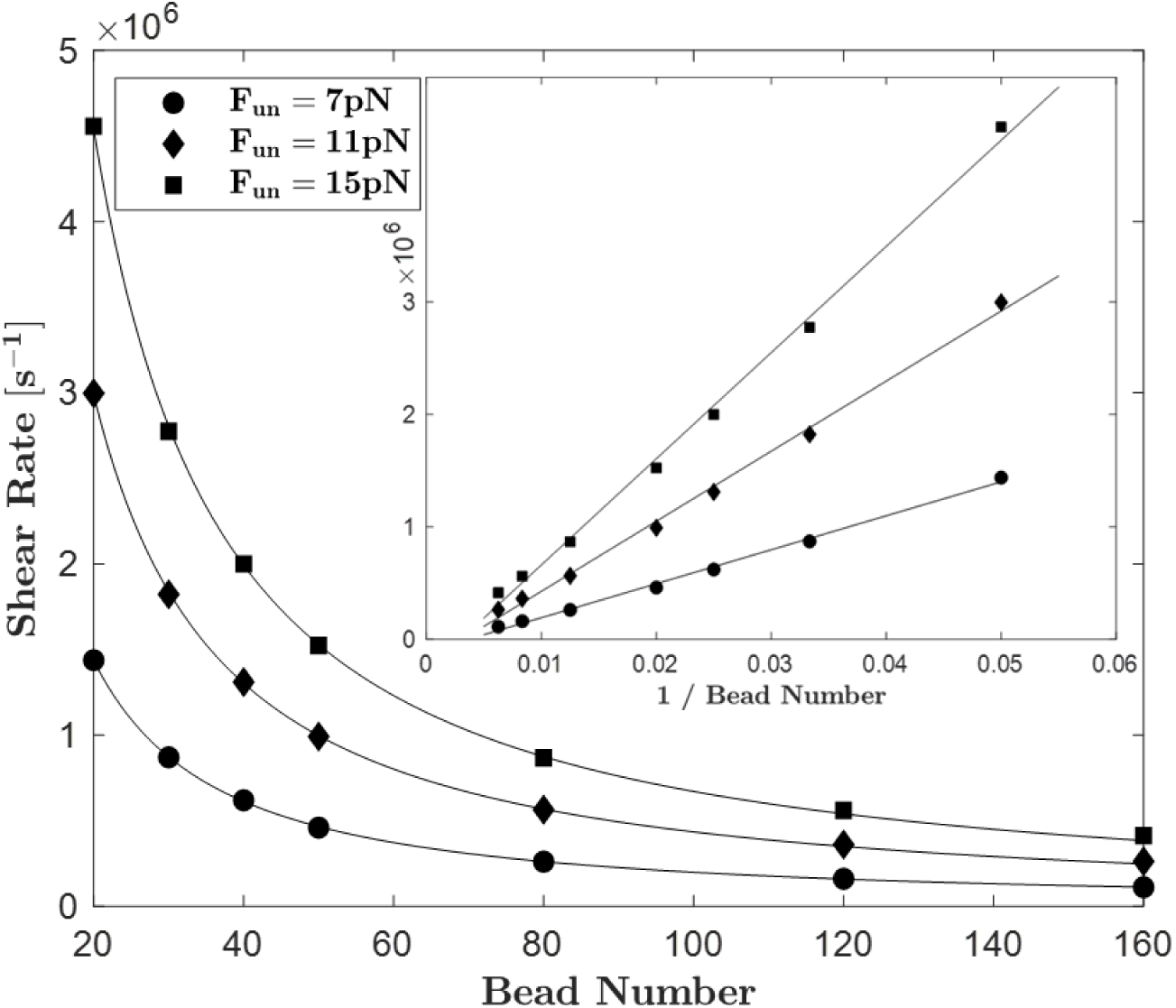
Threshold shear rates that enable A2 domain unfolding for at least the centermost monomer as a function of multimer length, as predicted by our analysis (see text). Three data sets and curve fits are shown that represent the threshold A2 domain unfolding forces of *F*_*UN*_ = 7,11, and 15 *pN*. Curve fits on the primary axes indicate threshold shear rate nearly depends on the inverse of chain length 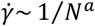, where the exponents are close to unity for all values of *F*_*UN*_. The curve fittings in the subfigure validate this inverse relation by showing a linear dependence of threshold shear rate on the inverse of chain length.

The dependencies in Fig. 8 are well fit by a power law, 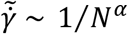, and the exponents corresponding to *F*_*UN*_ = 7,11, and 15 *pN* are *α* = 1.23,1.19, and 1.18, respectively. The subfigure shows a linear dependence of the same threshold shear rate data on the inverse of multimer length, 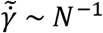. We have assumed that A2 domain unfolding is a flow-activated process, and observed that the threshold shear rate required to induce A2 domain unfolding inversely depends on multimer length, which is very similar to what a recent experimental work reported concerning another closely related flow-activated process for the A1 domain of vWF. Fu et al. studied flow-induced activation of the A1 domain, which enables binding to platelet GPIbα [15]. In that work, it was the assumption that A1 activation occurs at a threshold fluid shear stress. However, they showed that it is not shear stress that activates the A1 domain, but rather a threshold internal tensile force. They illustrated the dependence of shear stress (*σ*) on A1 domain activation as a function of chain length (*M*), which was well fit by *σ* = 1/*M*. The dependence characterizing this closely related flow-activated process is very similar to what we report for A2 domain unfolding in Fig. 8. Those authors offer strong support that a constant internal tensile force induces A1 domain activation, which is what we have assumed in this work for A2 domain activation. Accordingly, the trends we observe are in good agreement with experimental findings.

Extrapolation of the middle curve in Fig. 8 to physiological shear rates yields the multimer length for which A2 domain unfolding will, on average, readily occur in the bulk. At the physiological shear rate 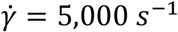, the relation predicts that a vWF multimer comprised of ~ 2,000 monomers would be readily cleaved for the unfolding threshold force of 11 *pN*. Other authors have made connection between simulation results and shear-induced vWF-related phenomenon by specification of a single physiological shear rate [10], [40]. It is important to make clear the distinction between macromolecular unraveling events examined by others and intra-monomer FENE spring force excursion events that we associate with A2 domain unfolding here. Fig. 1 clearly demonstrates that our model yields the proper macromolecular behavior with increasing shear, because growth in Rg (i.e. globule-stretch transition) begins around the Wi of unity for all multimer lengths. The large shear rates required to induce A2 domain unfolding may suggest that cleavage rarely occurs in the bulk [41] – perhaps A2 domain unfolding and subsequent scission occur more readily upon interaction with a surface or ancillary particle. In fact, other authors have shown that tethered vWF are much more susceptible to cleavage by ADAMTS13 [15]. Further, flow in the physical system is much more complex than a simple shear, and involves elongation that is much more effective at unfolding the A2 domain [42].

### d) A2 Domain Unfolding Analysis

The connection of A2 domain kinetics to our simulation model and results is made by defining an A2 domain unfolding event by any force excursion past the prescribed threshold unfolding force corresponding to 11 *pN*. The general assumption is that A2 domain unfolding is an activated process where transition from the folded to unfolded state is separated by a transition state energy barrier, which is lowered by the application of an external force. Unfolding events expose the hidden cleavage site and make the individual vWF monomer susceptible to scission. We can directly examine A2 domain unfolding probabilities, locations, and durations because we have explicitly considered the A2 domain in our monomer model.

The probability of an A2 domain to be unfolded and susceptible to cleavage is given by:

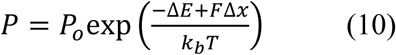

Others have examined hydrodynamic force-induced activation of collapsed biopolymers and the connection to the enzymatic cleavage of vWF by ADAMTS13 [9], [10]. Like these authors, it is assumed that A2 domain unfolding is the rate-limiting step in the scission of vWF by ADAMTS13 (i.e. that enzyme diffusion to the unfolded domain and the subsequent reaction are guaranteed). Accordingly, the frequency of A2 domain unfolding is given by 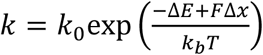. We analyze A2 domain unfolding probability in the following section, but we could have presented unfolding frequency instead.

Physiological vWF is subject to complex flow conditions that unravel the multimer (i.e. initiate the globule-stretch transition) and subsequently apply hydrodynamic force on the exposed A2 domain. Illustrated in Fig. 7, the average peak FENE force for only unraveled conformations varies linearly with shear rate. The average FENE force for only unraveled conformations also varies linearly with shear rate, however the ensemble average force does not because of the varying frequencies of A2 domain unfolding among chain lengths (not shown). We directly computed the probability of A2 domain unfolding by tallying the number of unfolded A2 domains (i.e. FENE spring force excursions past 11 *pN*) at each data collection step throughout the entire simulation for the 24-chain ensemble, then we normalized this quantity by the total number of possible unfolding events for all time, chains, and FENE springs.

The dependence of A2 domain unfolding probability on shear rate qualitatively exhibits exponential-like growth with increasing shear rate described by equation (10), because shearing provides the externally applied force required to lower the transition state energy barrier. Other authors examined vWF cleavage rates, which is effectively equivalent to A2 domain unfolding probability (in Fig. 9) in light of the assumption that it is the rate limiting step. Lippok et al. described their experimental vWF cleavage data by a phenomenological sigmoidal function of shear rate, and Radtke et al. prescribed this same dependence in their model for stochastic cleavage site opening, which is based on a theoretical two-state protein unfolding kinetics model [9], [10].

**Figure 9:**
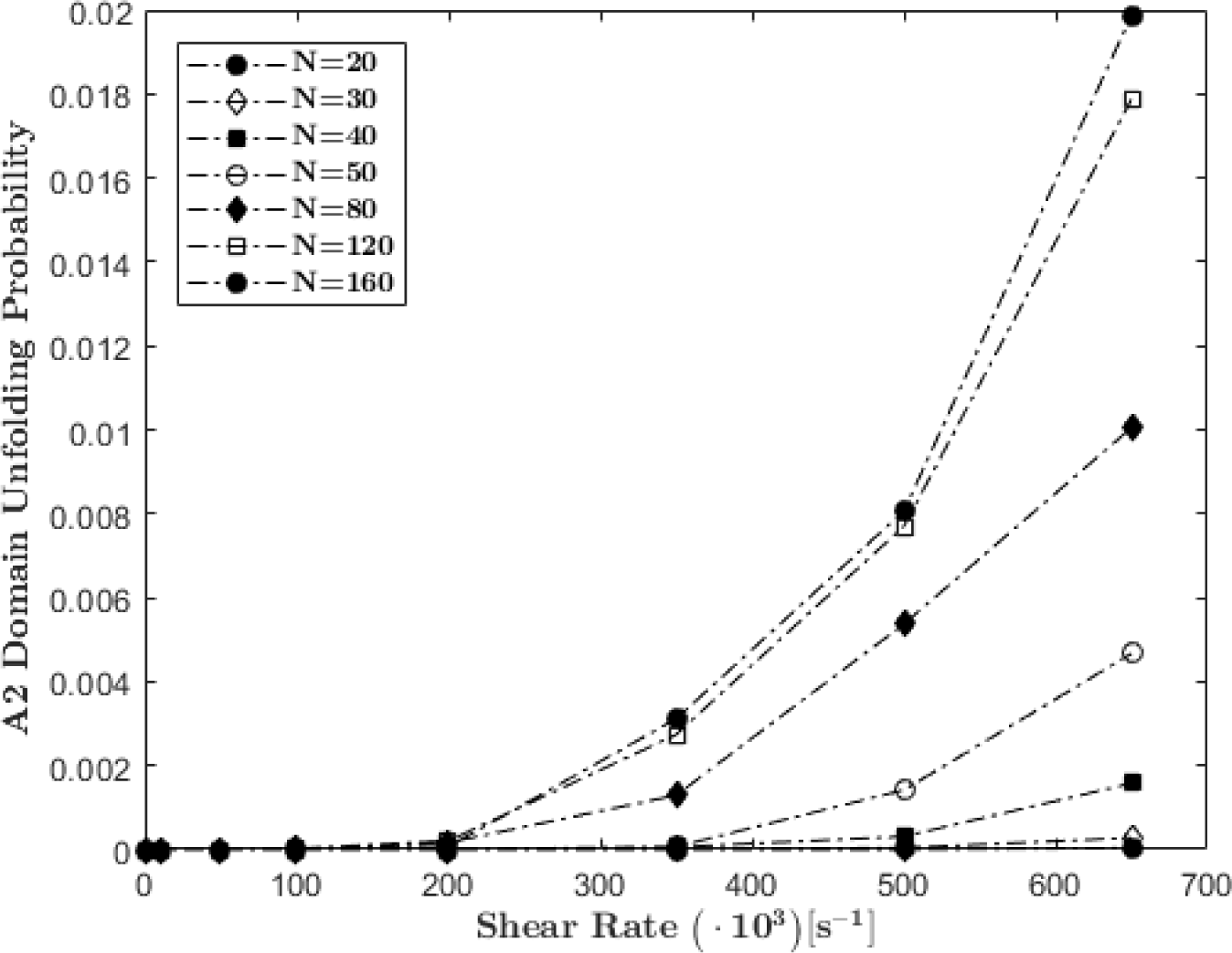
Probability of A2 domain unfolding as a function of shear rate for all considered multimer lengths (dashed lines are not curve fits).

Another method of varying the applied force is by examining various chain lengths at a sufficiently high but fixed shear rate. Force applied on a multimer by the fluid due to shearing at a fixed rate increases with multimer length because longer multimers have more drag forces acting upon them that must be internally balanced.

Figure 10 illustrates the probability of A2 domain unfolding as a function of bead number for several fixed shear rates. Interestingly, A2 domain unfolding probability does not vary according to what we assumed in equation (9), but rather exhibits a sigmodal behavior akin to what other authors have reported:

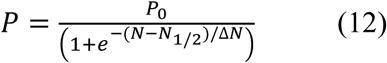

**Figure 10:**
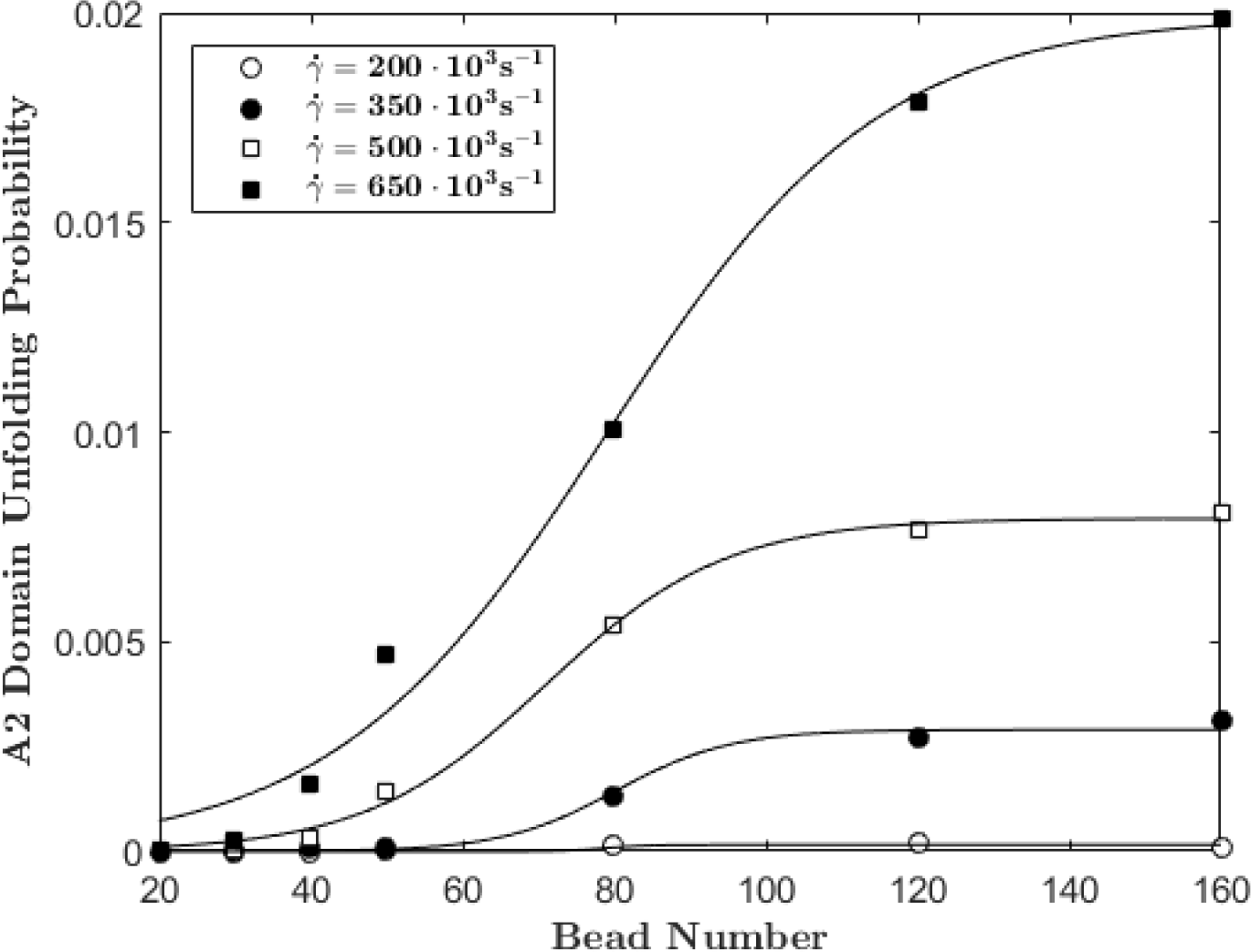
Probability of A2 domain unfolding as a function of bead number for several fixed shear rates.

*N*_1/2_, Δ*N*, and *P*_0_ are fitting parameters, where *N*_1/2_ represents the number of beads comprising a multimer for which the average probability of A2 domain unfolding is half of the maximum probability possible for the corresponding shear rate, *P*_0_. A2 unfolding probability monotonically increases with bead number. The probability increase is sharp for small chains lengths (*N* < 80), but increases more gradually for longer chain lengths (*N* > 80) and quickly plateaus. Longer multimers are able to resist the unravelling tendencies of shear flow because of increased HI shielding and macromolecular cohesive force. Hydrodynamic interactions induce shielding effects that mitigate the shearing force for downstream beads, which become increasingly dominant with chain length. Many additional pairwise additive Lennard-Jones attraction forces increase the multimers affinity for globule formation, where A2 domains are unlikely to unfold (see Fig. 3, 5, and 6). Clearly, vWF multimer length effects play a large role in determining A2 domain unfolding kinetics.

The force response behavior of A2 unfolding probability can be delineated according to introduction of external force by shearing or chain length variation. This A2 domain unfolding behavior appears to increase exponentially with shear rate and sigmoidally with multimer length. Explicitly modeling the A2 domain has allowed us to directly examine vWF A2 domain unfolding kinetics without enforcing any probabilistic conditions regarding the susceptibility of the cleavage site to ADAMTS13. A2 domain unfolding probability data in Fig. 9 and Fig. 10 did not discriminate according to the globular or unraveled state of the multimer, as we did for the force analysis discussed earlier.

It becomes difficult to explain the reported upper limit of in vivo vWF in light of the sigmodal dependence of A2 domain unfolding probability on multimer length, because multimers past a certain length are equally probable to transition from the folded to unfolded state. Accordingly, we assert that force excursions alone cannot be used as a hallmark for an unfolding event. This is supported by experimental data for pulling on vWF monomers where a rate effect has been observed for the most likely unfolding force [4]. Specifically, experiments have demonstrated that, for slower pull rates, A2 domains unfold, on average, at a lower pulling force. Furthermore, by applying pulling force at a relatively low magnitude and holding at that steady force for increasing time, experiments have shown that healthy, folded A2 domains will eventually unfold [4]. These observations indicate that it is not a force excursion alone that one should examine, but also the duration of the force excursions as well as the distribution of durations along the multimer.

Figure 11 illustrates the distribution of unfolding event durations for a multimer comprised of 80 beads (40-mer) subject to a shear rate of 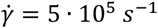. We simulated an ensemble of 24 non-interacting chains for thirty times longer than their relaxation time, *τ*, and results presented in Fig. 11 are per unit *τ*. The duration of unfolding events range from as little as one time step (~6.6 · 10^−4^ *μs*) to over one half-million time steps (~66 *μs*). We can see from Fig. 11 that the outermost ends of vWF multimers do not exhibit force excursions large enough to be considered for A2 domain unfolding; these end regions comprise 25% of the total contour length. The middle 75% of the multimer contour facilitates A2 domain unfolding, and the frequency of such events quickly decays with increasing unfolding duration. Examination of significant force excursion duration distributions in Fig. 11 suggests that A2 domains preferentially unfold near the multimer center, which is associated with A2 domain proteolysis.

**Figure 11:**
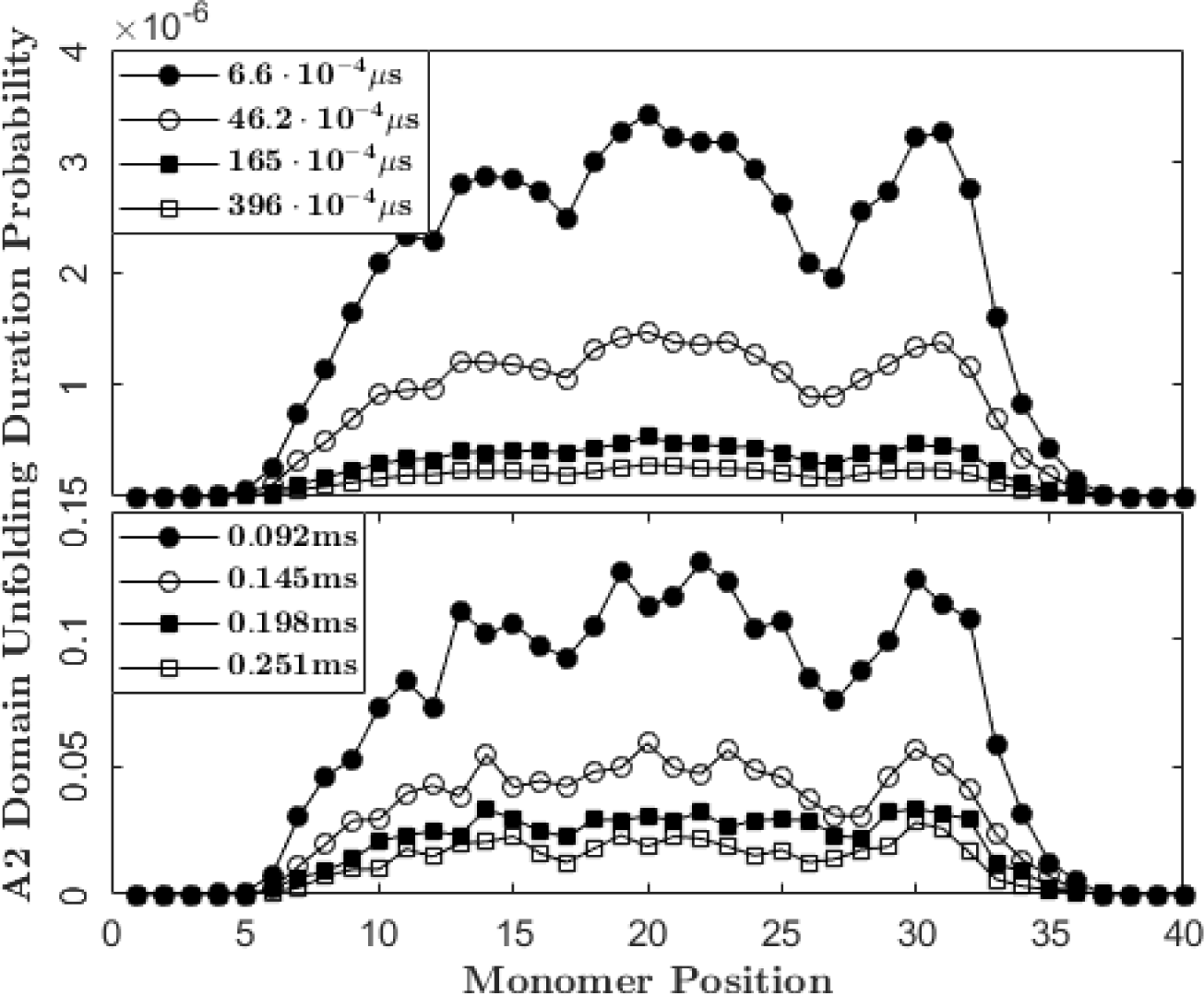
Probability of various A2 domain unfolding durations as a function of monomer position for a multimer of length N=80 (40-mer) at a shear rate of *γ* = 5 · 10^5^ *s*^−1^. Probabilities presented are per unit *τ*.

## CONCLUSION

We performed Brownian molecular dynamics simulations including hydrodynamic interactions to characterize the biomechanical response of vWF in bulk shear flow conditions. We explicitly modeled the A2 domain as a FENE spring, which we parameterized according to experimentally obtained A2 domain unfolding data. By granulating simulation results according to macromolecular conformation, it was observed that force monotonically increases with chain length only when shearing is actively unraveling the chain – this is not so for the ensemble average force. We suggest that A2 domain unfolding is most likely to occur for multimers that are unraveled or in the process of unraveling. Further, these multimers appear to be most susceptible to scission near the multimer center, where internal tensile force is highest. We exploited our explicit A2 domain model and directly assessed the susceptibility to scission for individual monomers based on FENE spring force excursions past the reported most probable A2 domain unfolding force of 11 *pN* [4]. From there, we observed that the shear rate required to induce A2 domain unfolding is inversely proportional to multimer length. Lastly, we examined A2 domain unfolding probability, which increases exponentially with shear rate and sigmoidally with multimer length. It will be of interest to our future work to determine how predictions made here change if one instead employs a model A2 domain that exhibits a finite minimum force for extension in association with an activated domain unfolding process. Our results help elucidate the internal force mechanisms at play in the platelet binding and scission processes unique to vWF. This further illustrates the power of increasing coarse-grain model complexity and capabilities – as motivated by experimental observations.

## ACKNOWLEDGMENTS

This work was supported by National Science Foundation grant DMS-1463234 and utilized the Extreme Science and Engineering Discovery Environment (XSEDE), which is supported by National Science Foundation Grant No. ACI-1548562 [43].

## AUTHOR CONTRIBUTIONS

All authors contributed to the design of this study. M.M. performed molecular dynamics simulations and data analysis. C.D. and W.W. aided in the interpretation of data. E.W., A.O., X.C., and X.Z. critically revised the manuscript and offered experimental/computational expertise.

